# Nanomotif: Leveraging DNA Methylation Motifs for Genome Recovery and Host Association of Plasmids in Metagenomes from Complex Microbial Communities

**DOI:** 10.1101/2024.04.29.591623

**Authors:** Søren Heidelbach, Sebastian Mølvang Dall, Jeppe Støtt Bøjer, Jacob Nissen, Lucas N.L. van der Maas, Mantas Sereika, Rasmus H. Kirkegaard, Sheila I. Jensen, Sabrina Just Kousgaard, Ole Thorlacius-Ussing, Katja Hose, Thomas Dyhre Nielsen, Mads Albertsen

## Abstract

DNA methylation is found across all domains of life but is a rarely used feature in recovery of metagenome-assembled genomes (MAGs). Recently, Oxford Nanopore introduced all context methylation detection models. We leveraged this to develop Nanomotif, which identifies and exploits methylation motifs for enhanced MAG recovery. We demonstrate how Nanomotif enables database-independent contamination removal from high-quality MAGs and host association of plasmids directly from Nanopore sequencing data in complex metagenomes.

## Main

In all domains of life, genomes are subjected to epigenetic modifications, which directly influences gene expression, replication, and repair processes^1^. In bacteria, the most common epigenetic modification is DNA methylation, which primarily acts as a host-defense mechanism against phages^1^. DNA methylation is facilitated by DNA methyltransferases (MTases), which recognizes specific DNA sequences, called motifs, and adds a methyl group to the DNA^1,2^. MTases often appear in restriction-modification systems, where a restriction enzyme recognizes a specific motif and cleaves the DNA if it lacks methylation. All DNA in the host must therefore have the correct methylation pattern for it to persist, including mobile genetic elements^2,3^. This evolutionary arms race has given rise to a great diversity of MTase recognition sequences^4^. Historically, DNA methylations have been identified using bisulfite conversions followed by short-read sequencing^1^. In recent years, Pacific Biosciences (PacBio) and Oxford Nanopore Technologies (ONT) have enabled direct detection of DNA methylations without the need for pre-treatment^5^. The most common methylations in bacteria are 5- methylcytosine (5mC), N6-methyladenine (6mA), and N4-methylcytosine (4mC). PacBio was first to demonstrate *de novo* detection of DNA methylation^5^, but currently has a low sensitivity for 5mC which requires a high sequencing coverage (250x)^6,7^. During 2023-24, ONT introduced all context methylation detection models making 4mC, 5mC and 6mA methylation calls readily available with high sensitivity (https://github.com/nanoporetech/dorado). Despite this, only few efforts have been made to utilize ONT methylation calls for methylation motif discovery in bacteria^8–10^, but none which scales or extends motif discovery to metagenome sequencing of microbial communities.

In metagenomics, DNA methylation motifs are directly applicable in binning by clustering contigs, assess contamination in bins, and associate mobile genetic elements to specific microbial hosts. Previous studies have utilized methylation motif information for metagenomic binning and association of plasmids^2^. However, these methodologies suffer from the low PacBio sensitivity for 5mC^2,11^ or require whole genome amplification for detection of motifs using ONT^12^. Building on the recent methylation calling capabilities of ONT sequencing, we developed Nanomotif, a fast, scalable, and sensitive tool for identification and utilization of methylation motifs in metagenomic samples. Nanomotif offers *de novo* methylated motif identification, metagenomic bin contamination detection, association of unbinned contigs to existing bins, and linking of restriction-modification systems to methylation motifs (Fig. 1A).

**Fig. 1:**
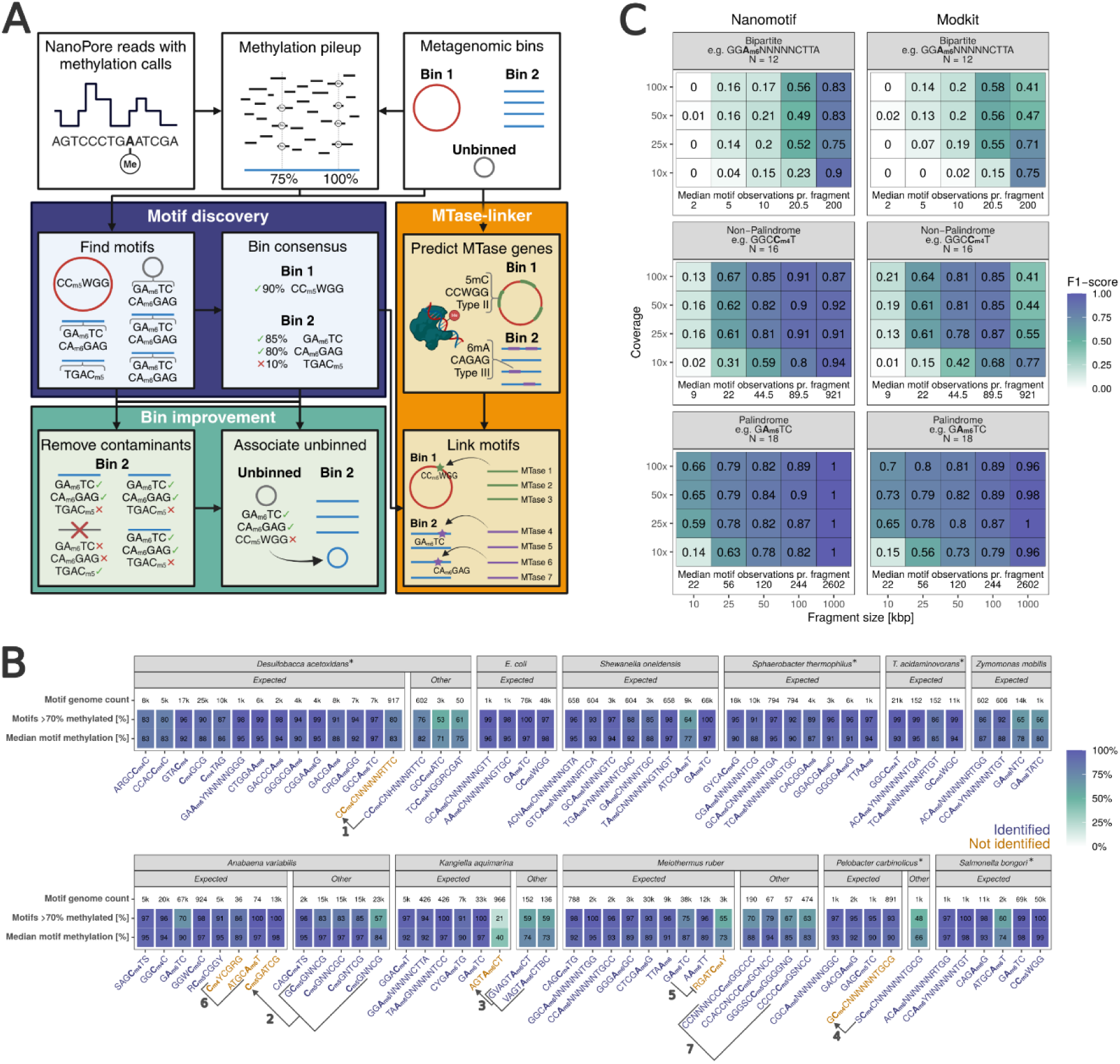
Nanomotif overview and benchmark. a, Overview of Nanomotif functionality. White boxes on the top row are required inputs for Nanomotif, colored boxes are Nanomotif modules. **b,** Motif identification using Nanomotif in 11 monocultures with 75 known motifs (*monocultures used only for testing). For each motif three values are shown: the number of occurrences in the genome, the percent of motif positions with methylation >70% and the median methylation of motif positions. Motifs are grouped by whether or not they were expected to be found or not (See supplementary data 1.a). Seven cases of faulty motif identification have been annotated 1-7. **c,** Benchmarking of palindrome, bipartite, and non- palindromic motif identification with Nanomotif and Modkit^7^ across different coverages and reference lengths. The number of expected motifs for each motif type is given in the header.

Nanomotif finds methylated motifs in individual contigs by first extracting windows of 20 bases upstream and downstream of highly methylated positions (>80% methylated). Motif candidates are then built iteratively using a beta-Bernoulli model, which evaluates whether the new candidate is more methylated relative to its originating candidate motif. The motif candidate search is directed using the KL-divergence from a non-methylated background, which rapidly guides Nanomotif through the motif search space, greatly decreasing search time compared to other algorithms (Supplementary note 1, Tab. S1).

We investigated a total of 28 monocultures, including 11 REBASE gold standard strains with known methylation motifs. The 11 monocultures with 75 expected methylation motifs were further split into 6 strains (29 motifs) for training and 5 strains (46 motifs) for testing. Firstly, the 29 motifs were used for parameter tuning (Fig. S1). We then benchmarked Nanomotif against Modkit^10^, MicrobeMod^8^ and MotifMaker^13^ on both training and test monocultures. Only Nanomotif and Modkit performed satisfactory and were therefore included in further benchmarks (Fig. S2 & S3). Nanomotif achieved a high recall rate and precision across all monocultures, identifying 68 out of the 75 expected motifs and 15 other motifs (Fig. 1B). Nine of the other motifs were closely related to a non-identified expected motif (Fig. 1B-#1-4). In *M. ruber*, RGAT**4mC**Y was missed, as it is a sub-motif of G**6mA**TC (Fig. 1B-#5). In *A. variabilis*, **4mC**YCGRG and ATGC**6mA**T were missed due to only 36 and 74 occurrences, respectively (Fig. 1B-#6). Lastly, four rare motifs were identified in *M. ruber* (57-474 counts) that likely represent noise due to increased 5mC false positive rate in high GC% organisms (Fig. 1B-#7 and Fig. S4).

To simulate metagenomic conditions, we further benchmarked motif identification by segmenting the test monoculture genomes to a varying number of fragment sizes and coverages (Fig. 1C). Nanomotif and Modkit perform similarly on palindromic motifs, which can be confidently identified in 25 kbp fragments even at 25x coverage. Palindromic motifs are generally shorter and therefore easier to detect because of their higher frequency and simplicity. For non-palindromic and bipartite motifs Nanomotif and Modkit have similar performance on 10-100 kbp, however on 1000 kbp fragments Modkit finds more false-positives leading to a drop in precision and hence F1-score (Fig. S5 & S6). Lastly, we benchmarked scalability, where Modkit used 23-40 times more total time compared to Nanomotif (Tab. S1). Modkit is limited to a single bin per instance, which requires all input files to be split into single bins beforehand (see Methods). The time spent on preprocessing input files was not included in the benchmark results.

A unique feature of Nanomotif is the scalability to complex metagenomic samples. We therefore used Nanomotif on five increasingly complex metagenomic samples (Fig. 2A).

**Fig. 2:**
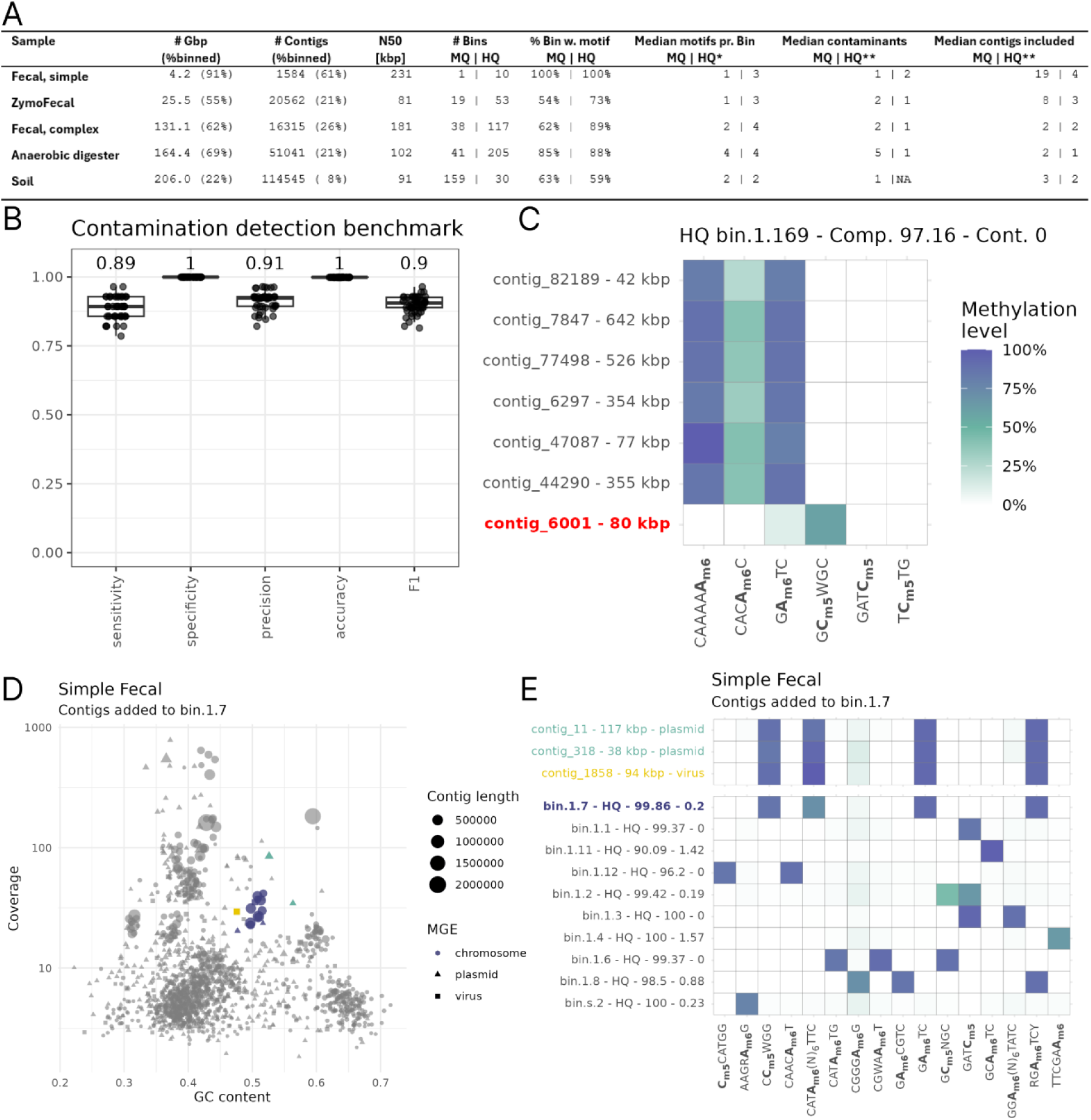
Nanomotif MAG contamination detection and association of mobile genetic elements. a, Sample stats from binning and the Nanomotif modules. Only values for bins above 10x coverage are reported. *The median is reported for bins with at least one motif identified. **The median is reported for bins where contaminants were removed or contigs included. **b,** Nanomotif detect_contamination benchmark metrics from 50 benchmark datasets. 28 monocultures at 40x coverage were fragmented into one 1600 kbp fragment and several 20 kbp fragments. One randomly selected fragment was randomly assigned to another bin in each benchmark dataset. **c,** Example of contamination removal (red) from a HQ MAG from an anaerobic digester. **d,** GC% and coverage of bin.1.7 in the simple fecal sample (blue). Three contigs, predicted as two plasmids and one virus, are assigned to the bin with the Nanomotif include_contigs module. **e,** Methylation profile of the HQ bins in the simple fecal sample and highlighted plasmid & viral contigs assigned to bin.1.7.

Across all metagenomic samples, at least one motif was identified within 87% of metagenome-assembled genomes (MAGs) above 10x coverage, and within these the median number of identified motifs was 3. This is more than previously reported in small-scale meta-epigenomic studies, which only identified methylation motifs in approximately 50% of MAGs using PacBio^14,15^.

Building on the motif discovery algorithm, we developed three modules for Nanomotif, which uses the motif methylation pattern; MAG contamination detection, inclusion of unbinned contigs, and linking of motifs to the responsible methyltransferases.

Current MAG contamination evaluation tools rely on lineage-specific markers derived from genome databases^16–18^, however, it is a difficult task as the databases are far from complete, and exceptions exist even within closely related organisms. To enable de novo contamination detection in MAGs, we leveraged Nanomotif to identify methylation motifs and then used ensemble clustering on the methylation pattern of bins (see methods). The 28 monocultures were used to benchmark the contamination module by fragmenting the monocultures into one 1600 kbp fragment and several 20 kbp fragments and then randomly assigning 20 kbp fragments to other bins (Fig. 2B, see methods). Nanomotif was able to achieve high sensitivity and precision with a mean of 89% and 91%, respectively. Most monocultures had near perfect contamination removal across all benchmarks (Fig. S7). We then applied the contamination detection module to the five real metagenomes of increasing complexity. The median number of contaminants in MAGs where at least one contaminant was detected was 1-2 for HQ MAGs. For example, bin 1.169 (HQ MAG) from the anaerobic digester (Fig. 2C) included contig_6001 (80 kbp) that completely lacked CAAAA**6mA** and G**6mA**TC methylation, despite the remaining bin being methylated at >75% in these motifs. In total, 196 contaminants were removed across 90 MAGs from the complex communities (Fig. S8). Each putative contaminant was manually reviewed, and in 84 out of 90 MAGs, the removal appeared accurate based on the methylation pattern, which matches the precision observed in the benchmark. This indicates a high potential for methylation to serve as a powerful post-binning cleanup, especially as this information is directly available for all new Nanopore sequencing projects.

The Nanomotif contig inclusion module assigns unbinned contigs to existing bins by training a linear discriminant analysis model, random forest, and k-neighbors classifier on the decontaminated bins (see methods). In case all three classifiers agree with a joint mean probability >0.8, the contig is assigned to the bin. Nanomotif achieved a high precision of 96% and moderate recall of 66% across the 28 fragmented monocultures described above (Fig. S9). In the five real complex metagenomic samples, the include module added a median of 1–4 contigs per bin for HQ MAGs. Associating mobile genetic elements with MAGs is of major importance as these can carry vital functionality^19^. This can be very difficult for traditional binners due to large variation in coverage or GC-content from the host but should be possible if a unique methylation motif signal is present. For example, using methylation motifs three contigs from the Simple Fecal sample were assigned to bin 1.7 and classified as two plasmids and a virus (Fig. 2d-e). It should be noted that the inclusion module is not a binner and assignments should be considered putative as methylation motif patterns can be shared across MAGs, which is also reflected in our efforts to prioritize precision over recall.

Restriction-modification (RM) systems are often substantial obstacles to genetic transformation, which pose a significant barrier for the implementation of novel bacteria as cell factories^20^. Circumventing these systems through RM system evasion or through heterologous expression of the methyltransferases in the cloning host (RM system mimicking) has shown to increase transformation efficiency significantly^20,21^. Therefore, we developed the Nanomotif MTase-linker module, which links methylation motifs to their corresponding MTase and, when present, their entire RM system (Fig. S10, supplementary note 2 & 3, and supplementary data 2). Across 11 monocultures, 52 orphan MTases and 29 RM-systems (exclusive type IV) were identified. 19 RM-systems were associated with an active methylation motif, and a total of 42 out of 71 detected motifs could be linked to a single MTase gene or RM-system with high confidence. Across 549 recovered HQ MAGs from five metagenomic samples, nanomotif identified 3,123 MTase genes, of which 1,297 were associated with RM-systems. For 76% of the detected motifs, at least one candidate MTase with matching methylation characteristics was identified within the same genome. Additionally, Nanomotif, successfully generated a high-confidence set of target motif annotations for 232 MTases. Hence, Nanomotif has the potential to drastically increase the number of putative links between motifs and MTase genes, thereby vastly improving the molecular toolbox and the RM-system databases.

With Nanomotif, *de novo* motif discovery is now seamlessly possible with standard Nanopore sequencing, even for short and low coverage contigs from complex metagenomes. We provide simple implementations that utilize these motifs for robust identification of putative contamination in MAGs, association of mobile genetic elements to hosts, and linking of motifs to restriction-modification systems. This greatly enhances the resolution of metagenomic investigations, opening the possibility of linking extrachromosomal DNA elements to the host.

## Data availability

Sequencing data generated during the current study is available in the European Nucleotide Archive (ENA) repository, under the accession number PRJEB74343. Assemblies, bins, and output from Nanomotif are available at https://doi.org/10.5281/zenodo.10964193.

## Code availability

Nanomotif is available at https://github.com/MicrobialDarkMatter/nanomotif. Code for reproducing figures and supplementary resources can be found at https://github.com/SorenHeidelbach/nanomotif-article.

## Acknowledgements

The study was funded by grants from VILLUM FONDEN (130690, 50093), the Poul Due Jensen Foundation (Microflora Danica) and the European Research Council (101078234). We further acknowledge the Novo Nordisk Foundation within the framework of the Fermentation-based Biomanufacturing Initiative (FBM), (Grant no. NNF17SA0031362), and the Novo Nordisk Foundation (Grant no. NNF20CC0035580).

## Ethics

The simple fecal sample was collected as part of a study registered at ClinicalTrials.gov (Trial number NCT04100291). The study adhered to the Good Clinical Practice requirements and the Revised Declaration of Helsinki. The participant provided signed written informed consent to participate and allowed for the sample to be used in scientific research. Consent could be withdrawn at any time during the study period. Conduction of the study was approved by the Regional Research Ethics Committee of Northern Jutland, Denmark (project number N- 20150021). The complex fecal sample was collected at Aalborg University with consent from the provider to be used in this study.

## Materials And Methods

### Sampling

*Escherichia coli* K-12 MG1655 (labcollection), *Meiothermus ruber* 21 (DSM 1279), and *Parageobacillus thermoglucosidasius* DSMc 2542 were grown overnight in LB, DSMZ 256 Thermus ruber medium, and SPY medium, respectively. Genomic DNA for *Desulfobacca acetoxidans* ASRB2 (DSM 11109), *Sphaerobacter thermophilus* S 6022 (DSM 20745), *Thermanaerovibrio acidaminovorans* Su883 (DSM 6589), *Kangiella aquimarina* SW-154 (DSM 16071), *Anabaena variabilis* PCC 7120 (DSM 107007), *Pelobacter carbinolicus* Bd1 (DSM 2380) and *Salmonella bongori* 1224.72 (DSM 13772) was ordered from Leibniz-Institute DSMZ. Raw pod5 files for *Shewanella oneidensis* MR-1*, Cellulophaga lytica* Cy l20, LIM -21 (DSM 7489), *Kangiella aquimarina SW-154* (DSM 16071)*, Zymomonas mobilis* subsp. pomaceae Barker I and the remaining 15 monocultures used for contamination and inclusion benchmarks were acquired from s3://cultivarium-sequencing/MICROBEMOD-DATA- NOV2023/pod5/^8^. ZymoHMW (D6322), ZymoOral (D6332), ZymoGut (D6331) and ZymoFecal (D6323) were ordered from ZymoBIOMICS. The simple fecal sample was collected at Aalborg University Hospital at the Department of Gastrointestinal Surgery as part of a clinical trial (ClinicalTrials.gov NCT04100291)^22^. The complex fecal sample was collected at Aalborg University with consent from the provider. Sampling of the anaerobic digester sludge has been described elsewhere^23^. Sampling of soil has been described elsewhere^24^.

### DNA Extraction

DNA from cell pellets of overnight grown cultures of *E. coli* K-12 MG1655 and *M. ruber* 21 was extracted with the PureLink Genomic DNA mini kit (Invitrogen, Thermo Fisher Scientific, USA) following manufacturer’s instructions with final elution in DNAse/RNAse free water. DNA from cell pellets of *P. thermoglucosidasius (*DSM 2542) was extracted with the MasterPure Gram positive DNA purification kit (Biosearch Technologies (Lucigen)), according to manufacturer’s instructions with a 60 min incubation step and final elution in DNAse/RNAse free water. DNA from ZymoOral (D6332), ZymoGut (D6331) and ZymoFecal (D6323) was extracted with the DNeasy PowerSoil Pro kit according to manufacturer’s instructions and suppliers suggestions. DNA from the simple fecal sample was extracted with the DNeasy PowerSoil Pro kit as described previously^25^. DNA from Complex fecal sample was extracted using DNeasy PowerSoil Pro kit according to manufacturer’s instructions. DNA was extracted from the anaerobic digester as described previously^23^.

### Sequencing

All samples were sequenced on the Promethion24 using the R10.4.1 nanopore. Anaerobic digester, complex fecal and ZymoFecal (D6323) were prepared with SQK-LSK114. Monocultures, ZymoOral (D6332), ZymoGut (D6331) were prepared with SQK-RBK114.24. ZymoHMW (D6322) was prepared with the SQK-NBD114-24 ligation kit. Sequencing of simple fecal is described elsewhere^22^. Sequencing of soil is described elsewhere^24^. All samples were basecalled with Dorado v0.8 using the dna_r10.4.1_e8.2_400bps_sup@v5.0.0 model and DNA methylation was called with the respective v2 methylation models for 4mC_5mC and 6mA.

### Assembly and binning

All samples were assembled and binned using the mmlong2-lite v1.1.0 pipeline available at^26^. Briefly, metaFlye (v2.9.4)^27^ is used for assembly and eukaryotic contigs are removed with Tiara (v1.0.3)^28^ before assembly coverage is calculated with read mapping via minimap2 (v2.28)^29^. Binning is performed iteratively as an ensemble using SemiBin2 (v2.1.0)^30^, MetaBat2 (v2.15)^31^, VAMB (v3.0.3)^32^, and COMEbin (v1.0.4)^33^ whereafter the best bin is chosen with DAS tool (v1.1.3)^34^. MAG quality was classified according to the MIMAG definition (Bowers et al., 2017). Completeness and contamination were evaluated with CheckM2 while rRNA and tRNA genes were found with barrnap (v0.9, https://github.com/tseemann/barrnap) and tRNAscan-SE (v2.0.16)^35^, respectively.

### Methylation pileup

Reads with methylation calls were mapped to the assembly using minimap2 v2.24^29^ using default settings. Nanopore’s modkit v0.4.0 (https://github.com/nanoporetech/modkit) was used to generate the methylation pileup from mapped reads using default settings.

### Motif identification

MicrobeMod v1.0.3 with default settings was used for all motif identification experiments. motifMaker (smartlink v13.1.0) with default settings was used for all motif identification experiments. Modkit pileup is not directly compatible with motifMaker and had to be converted to the same format as the output of ipdSummary. As the goal was a comparison of the motif identification algorithm, we extracted generally methylated positions (>70% methylated) and generated a GFF formatted file similar to the output of ipdSummary, marking all extracted positions with high Q-score and IPDRatio. Modkit v0.4.0 was used for all motif identification experiments. Default parameters were used for full genomes and scalability experiment. The setting --min-sites 20 was used for benchmark with lowered coverage and fragmentation of reference. This was done for fair comparison to Nanomotif minimum motif count of 20.

Nanomotif v0.5.0 was used for all experiments. Nanomotif motif discovery algorithm has two main submodules, “find-motifs” and “bin-consensus”. All subcommands are gathered in a parent command “motif_discovery”, which was executed with the following arguments for all samples: threshold_methylation_confident=0.8, threshold_methylation_general=0.7, search_frame_size=41, threshold_valid_coverage=5, minimum_kl_divergence=0.05, min_motif_score=0.2. “find-motifs” identifies motifs in contigs, referred to as directly identified motifs. This is done using a greedy search and candidates are selected based on a Beta-Bernoulli model, where each motif occurrence is Bernoulli trial, being a success if the fraction of methylation of reads at the position is above a predefined threshold (default 0.70). The Beta-Bernoulli was chosen in order to include uncertainty in the motif scoring process, instead of a point estimate. The exact steps performed for motif identification is outlined in the pseudo code below. For full details see supplementary note 1.

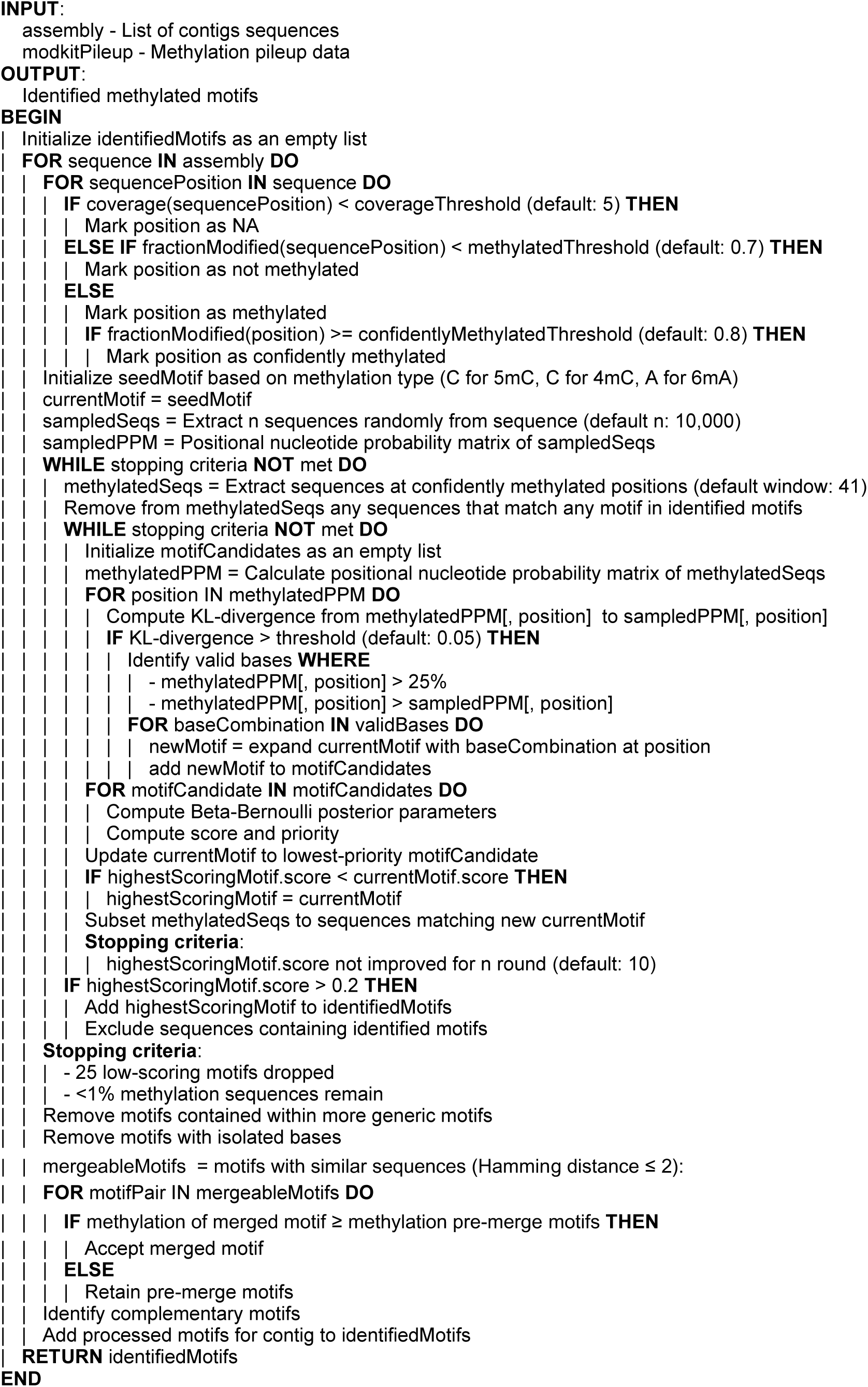

After identification of motifs in each contig “bin-consensus” evaluates the full set of identified motifs in the contigs belonging to the bin. It performs motif merging like the post processing steps in the motif identification algorithm, but for the bin motif set. Then motifs which are not methylated more than 25% methylated in 75% of the bin are removed.

### Motif Identification Benchmark

Motif identification was benchmarked using motifs identified in 11 monocultures with known methylated motifs. Full set of expected motif and motif evidence is available in supplementary data 1.a. Six monocultures were used for parameter justification of the “find-motifs” algorithm; *Shewanella oneidensis* MR-1*, Kangiella aquimarina* DSM 16071*, Anabaena variabilis* ATCC 27893*, Escherichia coli* K-12 substr. MG1655*, Meiothermus ruber* DSM 1279*, Zymomonas mobilis* subsp. pomaceae. A grid search was performed over the three most important parameters, to justify the final parameter settings (Fig. S1). The algorithm performance is stable across the selected parameters, and the final set parameters were chosen to increase sensitivity in metagenomic settings. Five monocultures were used for testing; *Desulfobacca acetoxidans* DSM 11109*, Salmonella bongori* NCTC 12419, *Sphaerobacter thermophilus* DSM 20745, *Pelobacter carbinolicus, Thermanaerovibrio acidaminovorans* DSM 6589. Testing monocultures were not seen during tuning.

Benchmark metrics were calculated by comparing identified motif with expected motif in monocultures. If an identified motif matches an expected motif exactly, it counts as a true positive. False positives are counted as motifs not in the expected motif set. False negatives are counted as motifs in the expected set which are not identified. Both forward and reverse complement are counted as a motif for motifs which are not palindromic. For benchmarks, precision, recall and F1-score is reported.

Reduced information benchmarking was performed across two parameters; read coverage (10, 25, 50, 100x) and contig size (10, 25, 50, 100, 1000 kbp). Read coverage affects false positive and false negative in calling of generally methylated positions, as lower coverage is more sensitive to non-systematic false positive and false negative calls at the reads level. Lower coverage was achieved using Rasusa^28^ by subsetting the total length of reads to a multiple of the assembly length of the respective benchmarking organisms. As contig size is proportional with motif occurrences, smaller contigs will have fewer motif observations, thereby less information for motif identification. Differing contig sizes were created by chunking the reference genome of the monoculture to fix sized windows using SeqKit2 (v2.5.1) ^36^ (windows were not allowed to overlap). Up to 20 chunks from the procedure for each monoculture were included in the benchmark. To fairly compare across fragmentation lengths, we reduced the minimum required motif observations for modkit to 20, the same requirements Nanomotif utilises for motif identification in contigs.

To benchmark execution time of Modkit on metagenomic samples, we split the pileup file into separate files, each containing the information related to the contigs of a single bin. Then the assembly was split into fasta files, each containing the contig sequences related to a single bin. Execution time for preprocessing of files was not included in the reported run time or CPU hours. To identify motifs in a bin, we executed Modkit on the files corresponding to a single bin. Modkit was executed with a single thread, and parallelly executed in 144 instances.

### Contamination detection

Contamination is evaluated using “nanomotif detect_contamination” which uses ensemble clustering. In case all four clustering methods, HDBSCAN, Gaussian Mixture Model, Agglomerative Clustering, and spectral clustering marks a contig as an outlier, the contig is marked as contamination.

Firstly, motif methylation is calculated as follows: The mean read methylation for each motif observation is calculated for motif observations with at least 3 read mappings. The median value of these is then reported as the motif methylation. This methylation value was more robust for smaller contigs compared to the mean of means. Before clustering, motif methylation is filtered for each contig if the product of number of motif observations and the mean read coverage is less than 24. This way the methylation value of a contig with a one motif observation is trusted if the read coverage is at least 24 or the contig has at least 8 motif observations. After filtering, contigs with missing motif values are imputed with the bin mean and PCA is performed to reduce dimensions while retaining 90% of variance. Contigs are then clustered with HDBSCAN (min_samples=3, min_cluster_size=2), Gaussian mixture model (n_components = n_bins, covariance_type=”full”), Agglomerative clustering (n_clusters = n_bins, affinity = “nearest_neighbors”) and spectral clustering (n_clusters = n_bins). For each clustering method, the bin cluster is the cluster with the largest fraction of the bin length which must constitute at least 85% of the total bin length. In case all methods agree a contig does not belong to the bin, the contig is flagged as a contaminant. Contamination contigs are then assigned to the “unbinned” pool.

### Include contigs

The “nanomotif include_contigs” module will attempt to assign unbinned contigs to the bin after decontamination. The include_contigs module uses an ensemble of supervised machine learning techniques; random forest (n_estimators=100), linear discriminant analysis (solver = ”svd”), and k-nearest neighbors classifier (n_neighbors = 3) classifier to assign unbinned contigs to bins. Firstly, the three classifiers are trained on the binned contigs after dimensionality reduction (see contamination detection). Missing motif observations in unbinned contigs are then imputed with a pseudo value randomly chosen between 0 and 0.15, and projected into the lower dimensional space using the binned conversion matrix. A contig is assigned to a bin if all three classifiers agree and the mean probability of the three classifiers is above 0.80.

### Contamination and inclusion benchmark

A synthetic benchmark dataset was constructed for developing and evaluating the contamination and inclusion module. 28 monocultures (see Fig S6) were sampled to 40x coverage and fragmented into 20 kbp fragments. For each fragmented monoculture, a single long 1600 kbp contig was reconstructed by stitching together 80 consecutive 20 kbp fragments, while the remaining fragments were retained unaltered. Nanomotif were then used to find motifs anew and contamination and inclusion were evaluated using the found motifs. To evaluate the contamination module 50 datasets were created where one randomly chosen contig from each monoculture was randomly added to another. For the inclusion benchmark 50 datasets were created where a random contig was removed from each monoculture. The 1600 kbp contig was not shuffled or removed.

### MTase-Linker

The Nanomotif MTase-linker module initially uses Prodigal v.2.6.3^37^ for protein-coding gene prediction (default settings) followed by DefenseFinder v1.2.0^38^ to predict MTases and related RM-system genes. The output file defense_finder_hmmer.tsv is filtered for all RM-related MTase hits. When a single gene has several model hits, the model that yields the highest score is selected. The output file defense_finder_systems.tsv is used to determine whether the identified MTase hit is part of a complete defense system. MTase hits associated with non-methylation-mediated defense systems are excluded. Additionally, RM type IIG MTase hits not identified by DefenseFinder as part of a complete RM system are also removed.

Using hmmer (with parameter –cut_ga) the predicted MTase protein sequences are queried against a set of hidden markov models (PF01555.22, PF02384.20, PF12161.12, PF05869.15, PF02086.19, PF07669.15, PF13651.10, PF00145.21) from the PFAM database^39^, to predict the modification type (5mC or 6mA/4mC). Furthermore, to infer the probable target recognition motif, the MTase protein sequences are queried using BLASTP (Blast v.2.14.1) against a custom database of methyltransferases with known target recognition motif from REbase^40^. We employ a threshold of 80% sequence identity and 80% query coverage to confidently predict the target recognition motif. Lastly, the RM system, RM sub-type, mod-type, and predicted motif information for each methyltransferase gene are used to link methylation motifs to the genes. The pipeline identifies high confidence MTase-motif matches, labeled as “linked”, through either a precise match between the predicted motif and the detected motif or when a single gene and a single motif share a similar combination of methylation features, which are unique within a MAG. When a high confidence match cannot be elucidated, the MTase-Motif- linker assigns feasible candidate genes, with the corresponding motif type and modification type, for each motif.

### MGE classification

Contig were labeled as Mobile genetic elements (MGE) when classified as a viral or plasmid by geNomad (v1.7.0) with a score >0.75, had a mean coverage above 10x and a minimum length of 10kbp.

---

**Fig. S1:**
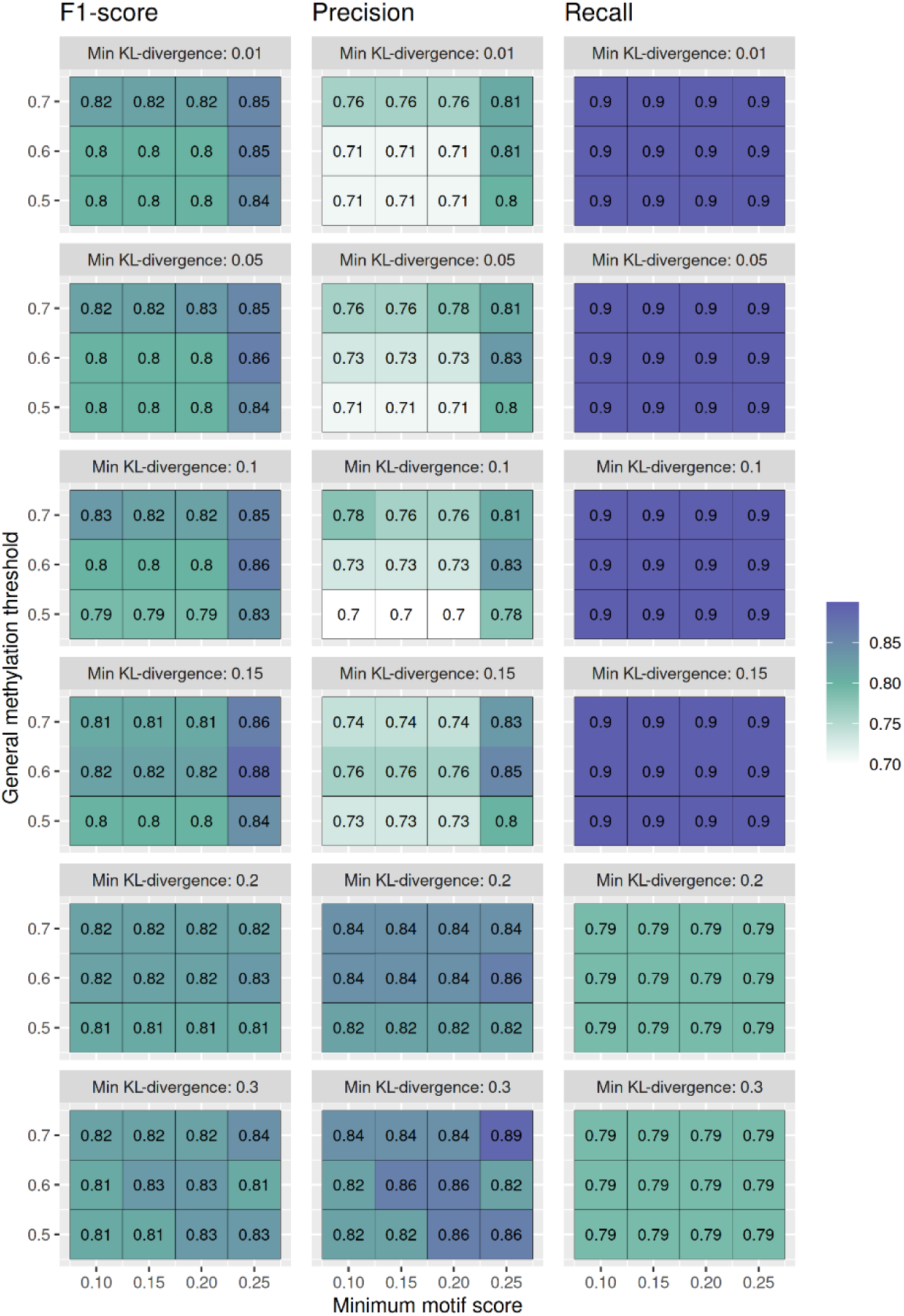
Parameter sweep over minimum KL-divergence, minimum motif score, and general methylation threshold used the “motif_identification” module. The parameter sweep was performed with expected motifs of the six training monocultures. Recall of expected motifs is primarily affected by the minimum KL-divergence parameter, whereas precision is reversely affected by the minimum KL-divergence. Precision increases with higher general methylation threshold and higher minimum motif score. Motif identification is generally stable across parameters.

**Fig. S2:**
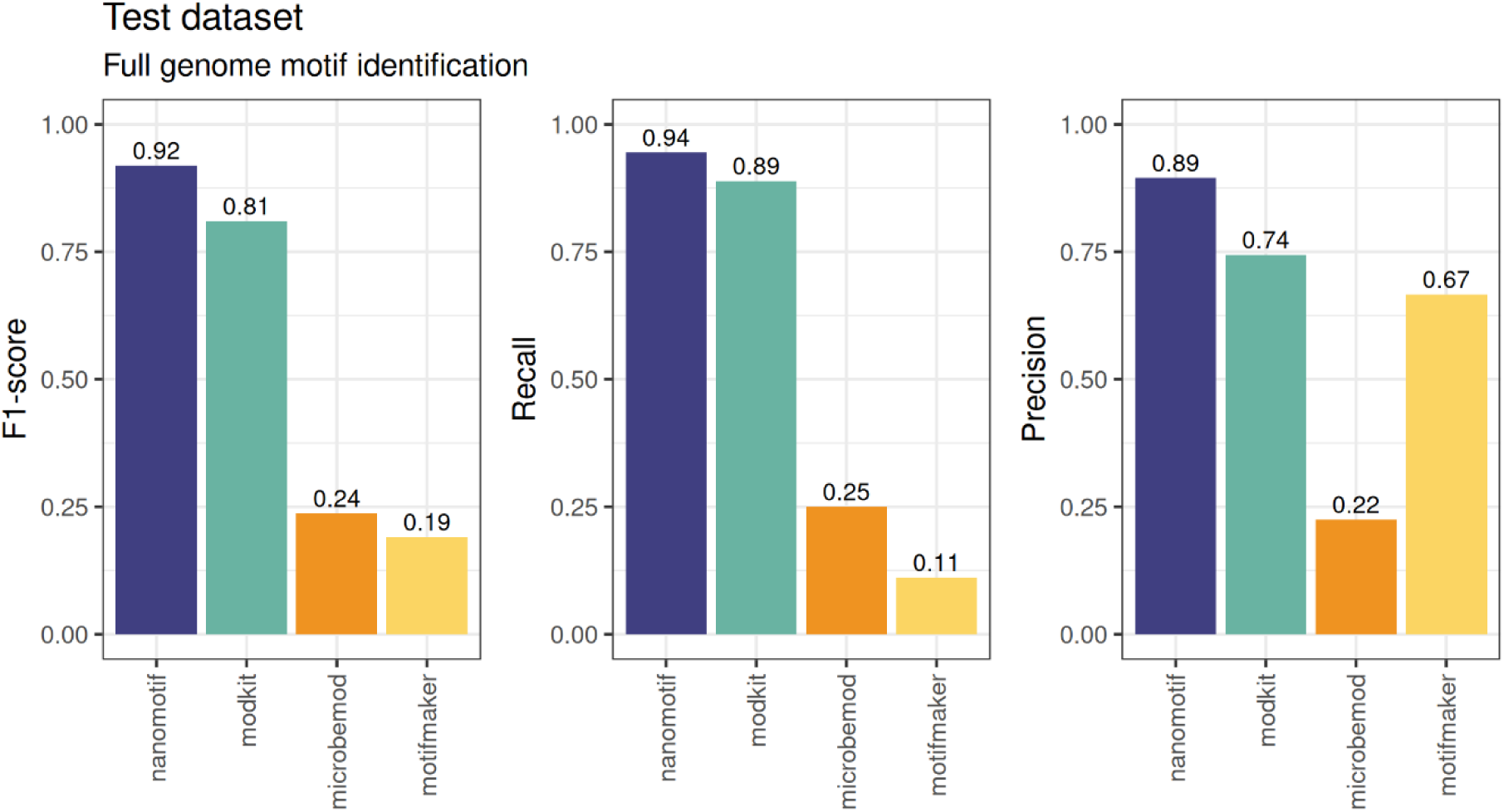
Full genome motif identification benchmark on 46 expected motifs from the five test monocultures.

**Fig. S3:**
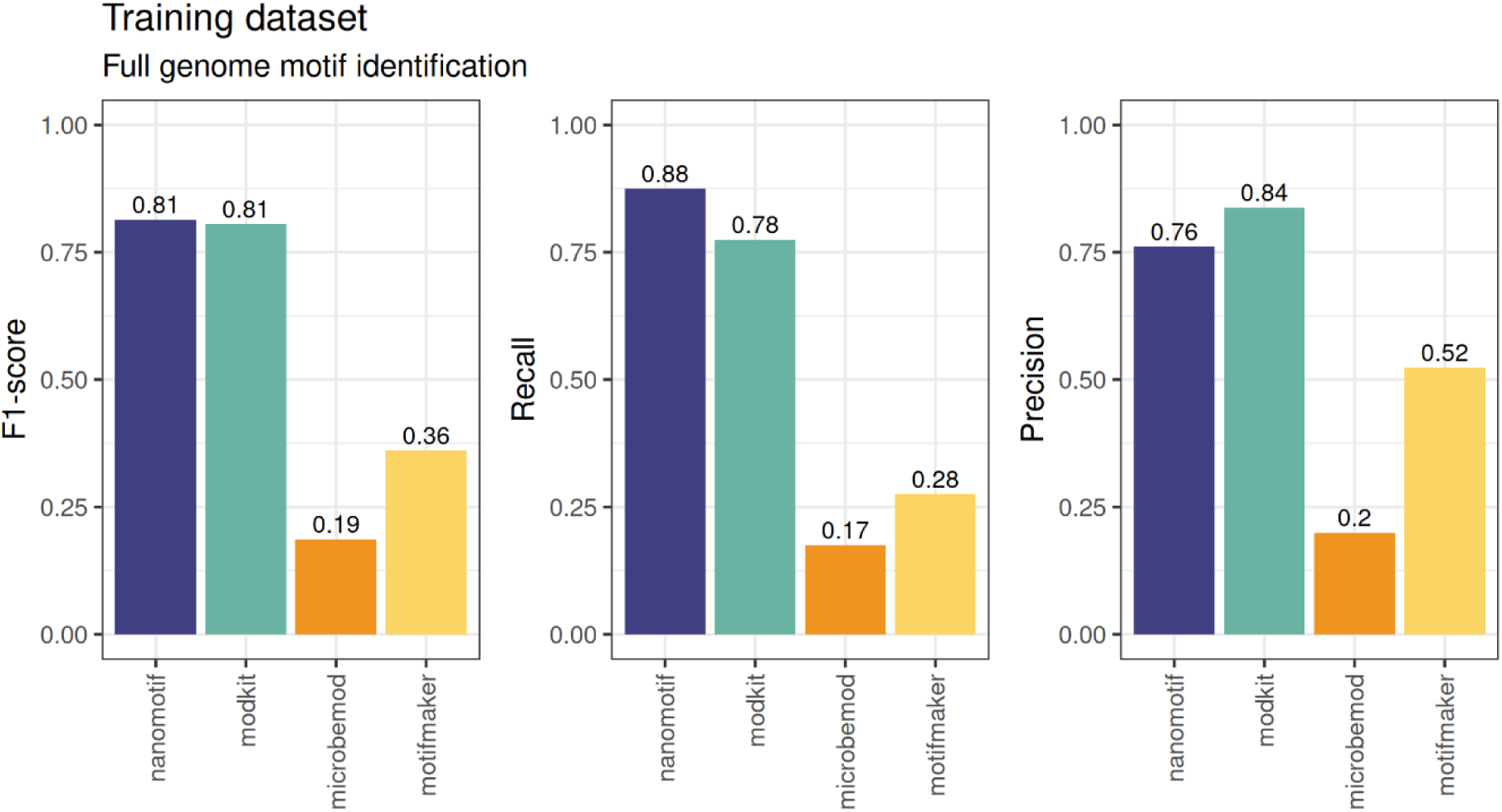
Full genome motif identification benchmark on 29 expected motif from the six training monocultures.

**Fig. S4:**
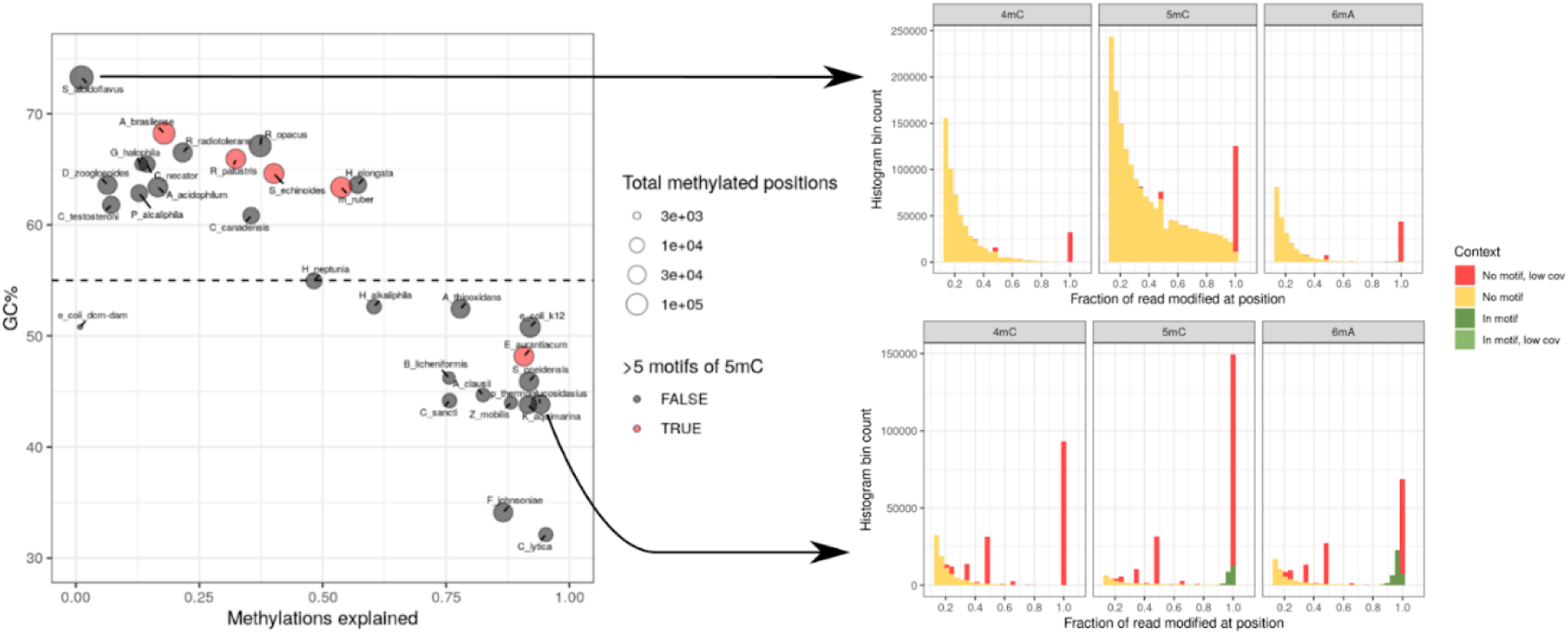
High 5mC false positives in high GC% species. Left panel; On the x-axis is the fraction of methylated positions occurring explained by a motif and the GC% of the organism’s genome. As the GC% increases, the degree of explained methylated positions decreases. Additionally, large degrees of false positive 5mC motif are identified in the high GC% organisms. Right panels; two example organismns, 1) *S. albidoflavus*, with a GC% of 74%, where 5mC has a continuous drop off in fraction of read modified at C positions, while none of these positions are explained by a 5mC motif. 2) *P. thermoglucosidasius*, with a GC% of 44%, where 5mC distribution is split into two groups; a low fraction group with no motif associated and a high fraction group, all explained by a motif.

**Fig. S5:**
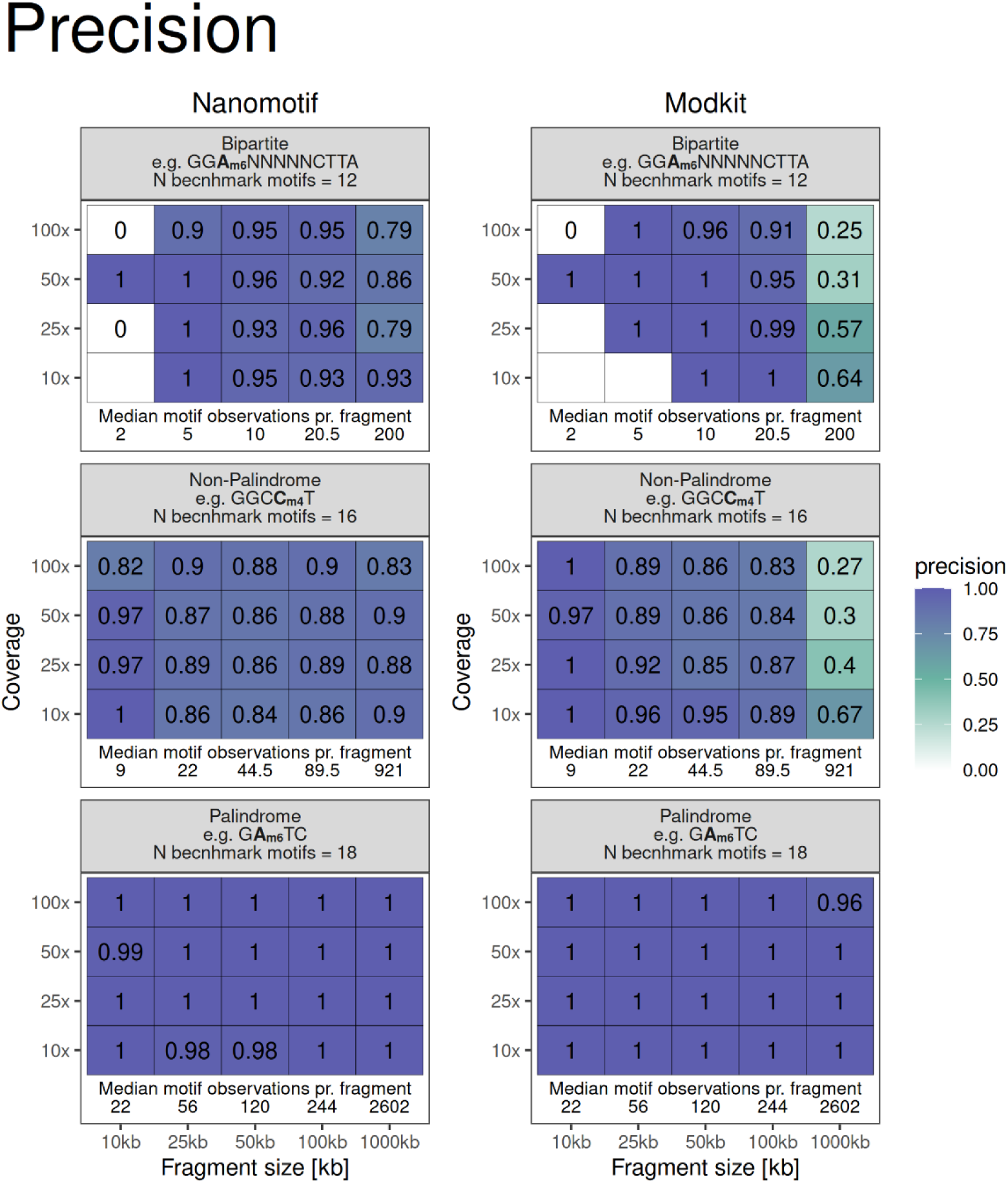
Precision of benchmark presented in Fig. 1C.

**Fig. S6:**
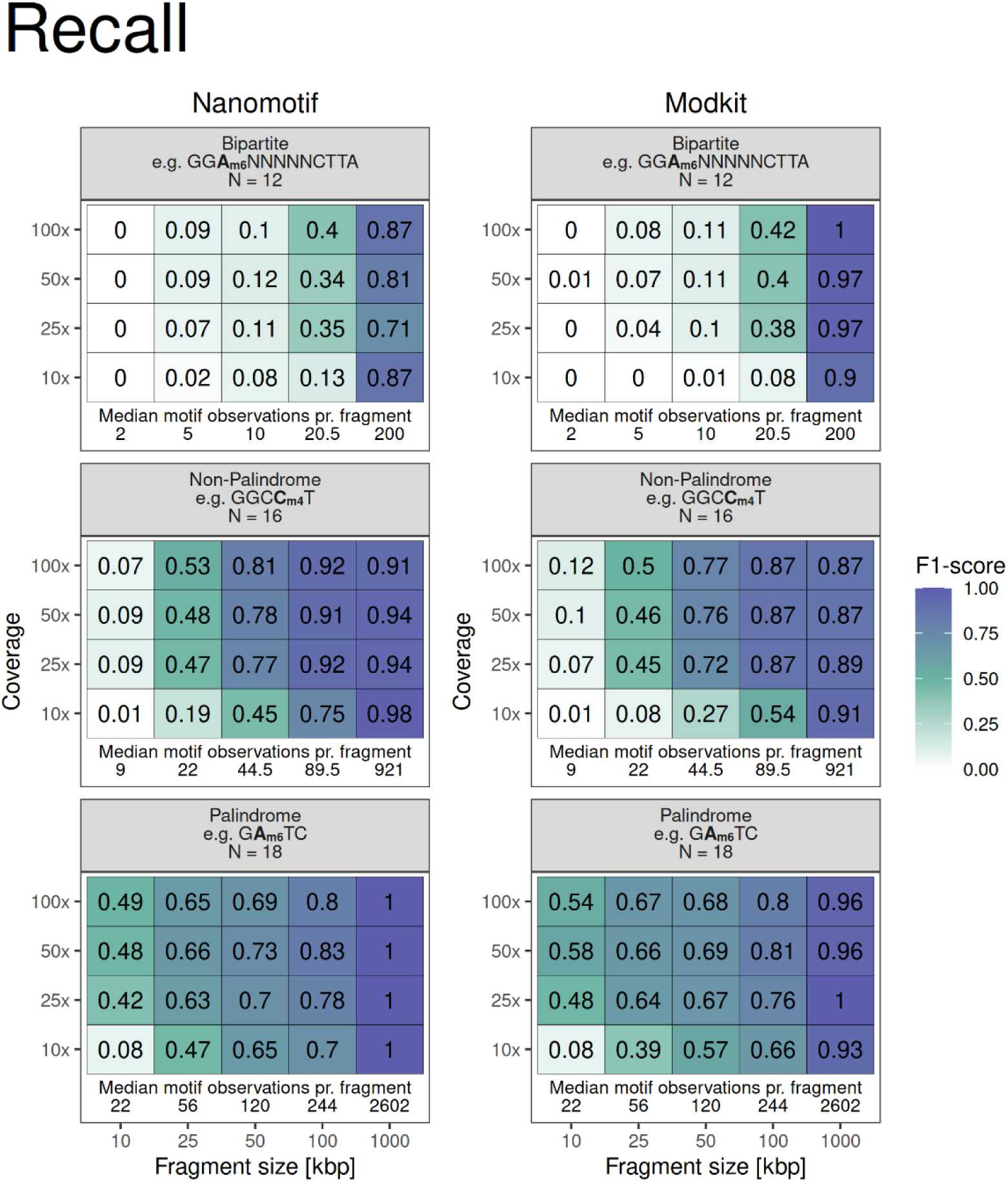
Recall of benchmark presented in Fig. 1C.

**Fig. S7:**
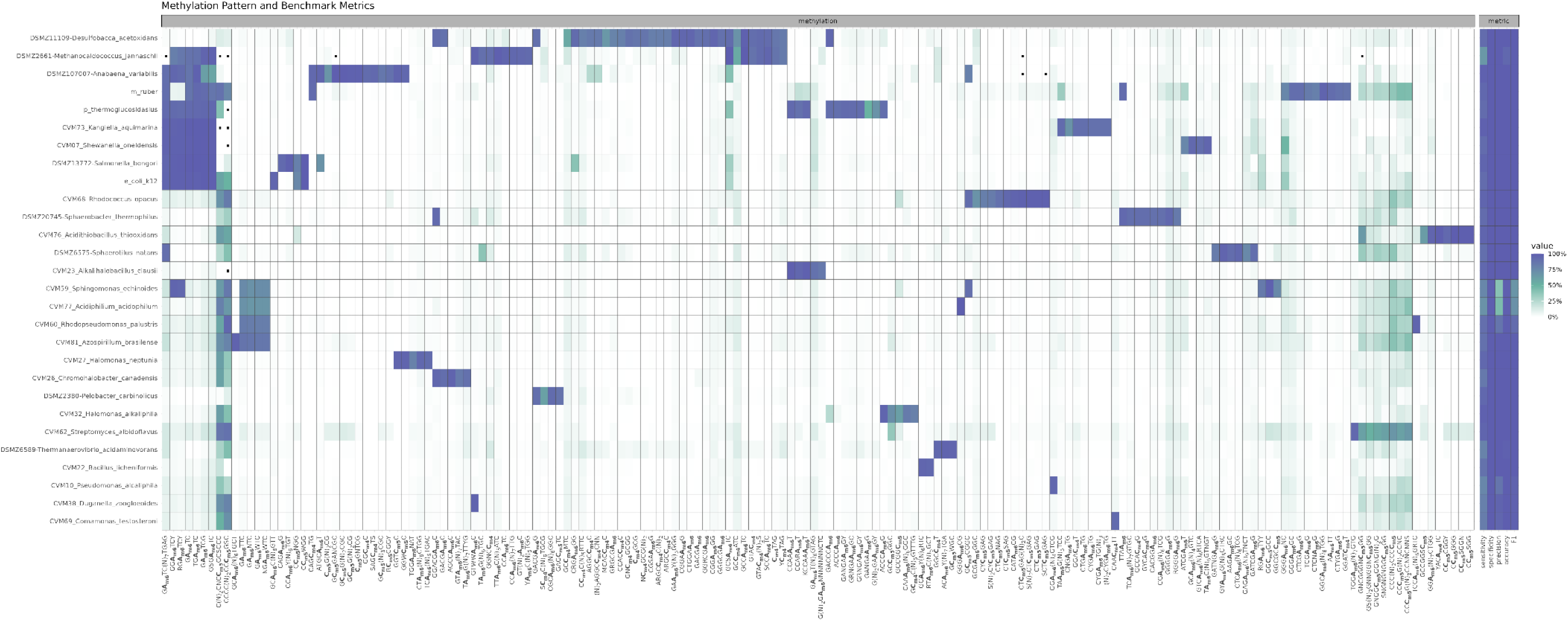
Methylation pattern of monocultures included in the benchmark dataset along with mean sensitivity, specificity, precision, accuracy, and F1 score across the 50 datasets where a contig from each bin has been assigned to another.

**Fig. S8:**
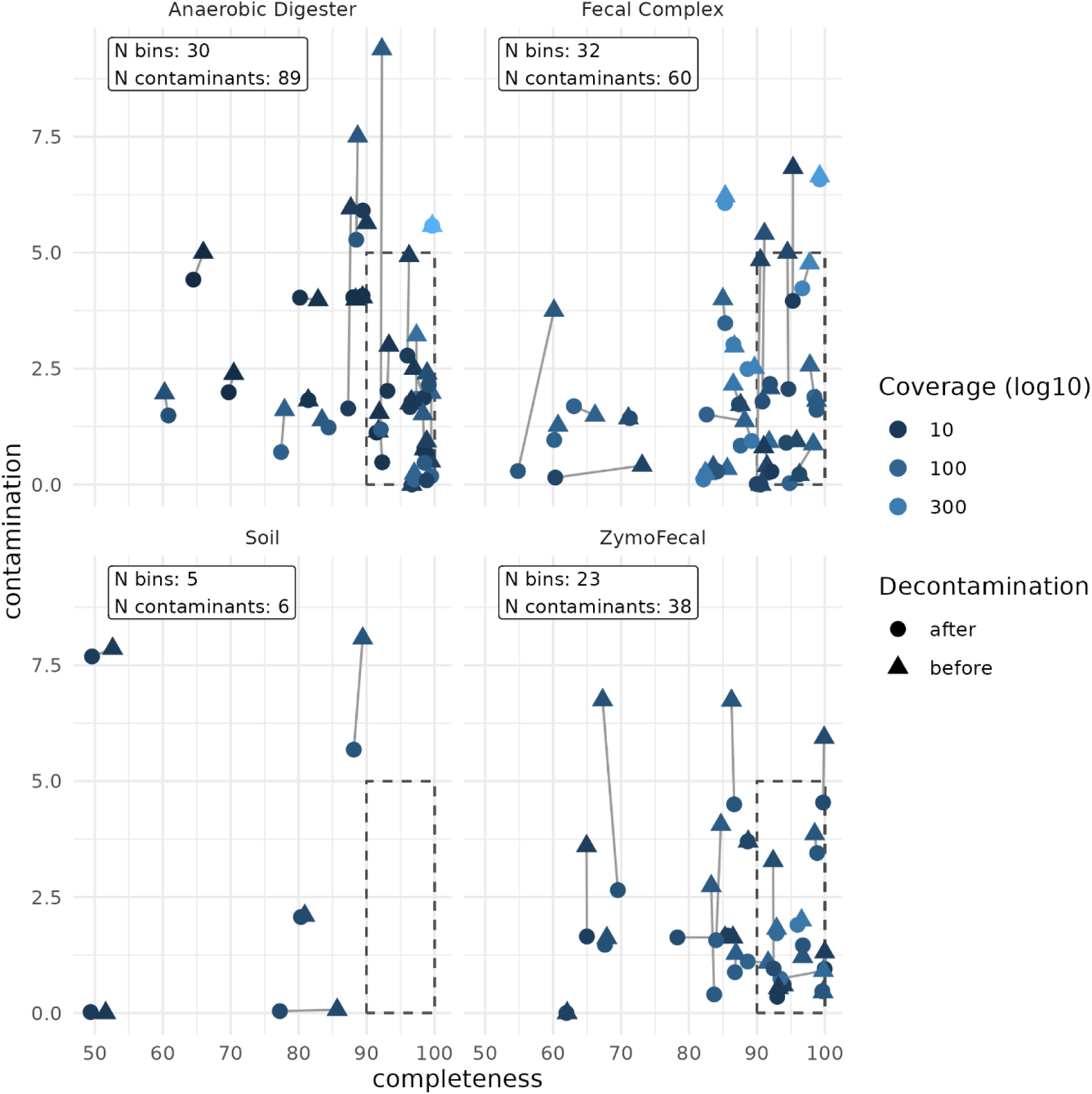
Completeness and contamination of MQ and HQ bins before and after removal of putative contamination. Dashed boxes mark completeness >= 90%, contamination <5%. One contaminant was removed in Fecal Simple but is not shown.

**Fig. S9:**
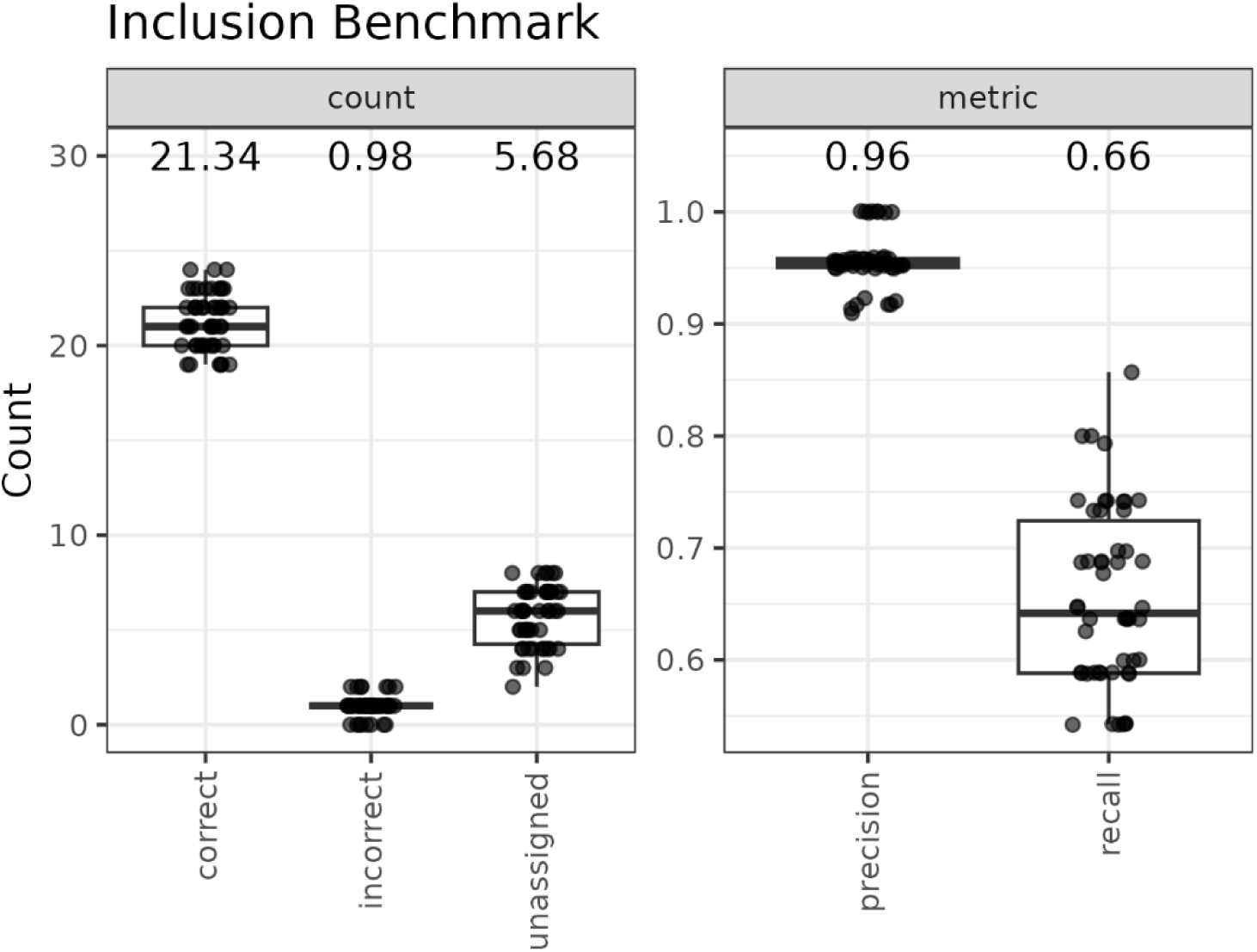
Number of correct, incorrect and unassigned classification from nanomotif include module using the monoculture benchmark.

**Fig. S10:**
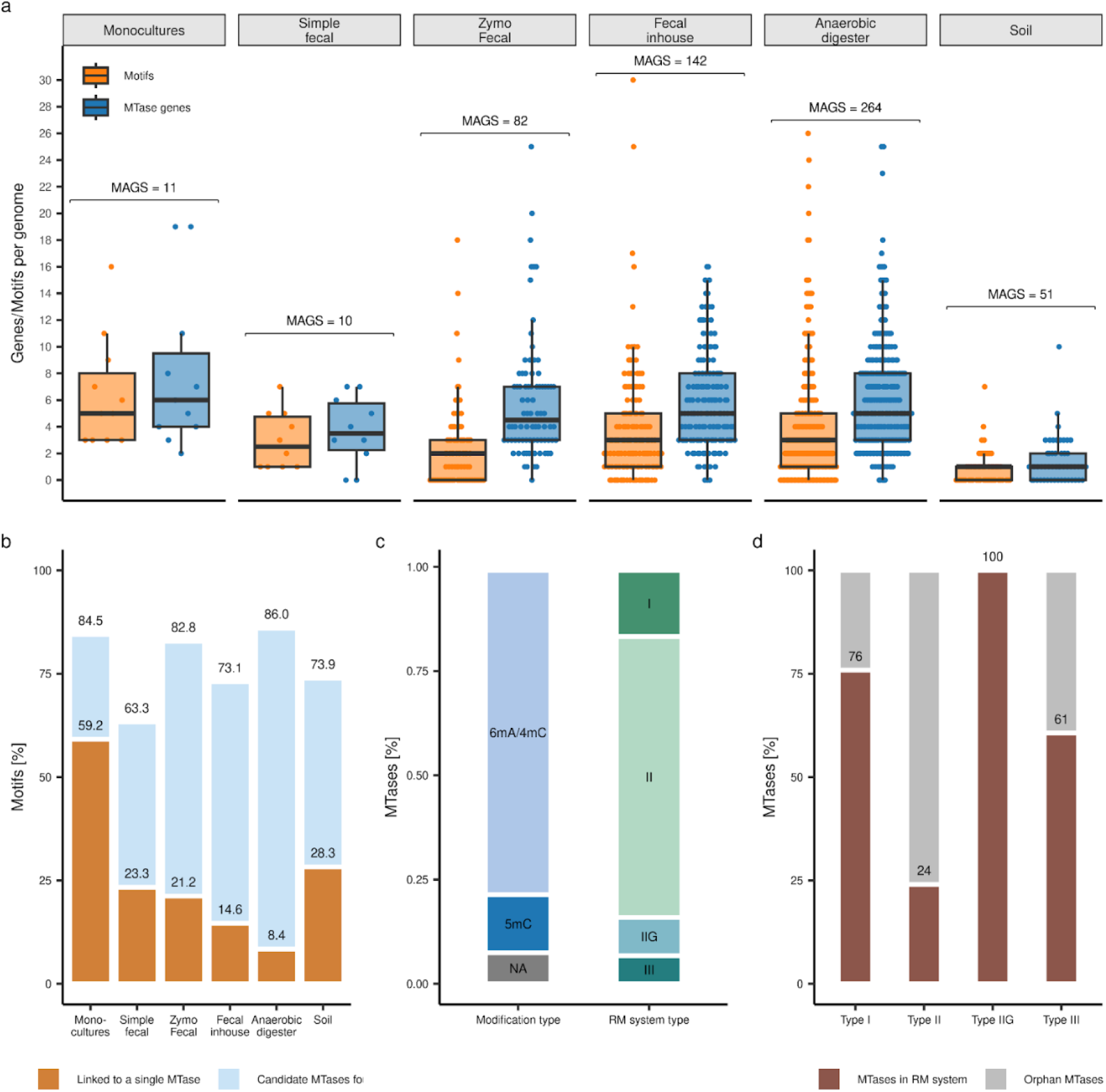
Summary of MTase annotations and motif assignments in 11 monocultures and recovered HQ MAGs from five metagenomes. **a**, Distribution of detected motifs and annotated MTase genes per. genome. **b**, Percentage of motifs with a high confidence linked to an annotated MTase gene (orange), or motifs for which one or multiple candidate genes have been found (blue). **c**, Breakdown of annotated MTases by modification type and RM-system type. NA indicates unidentified modification type. **d**, Proportion of annotated MTase involved in RM-systems.

**Tab. S1:** Benchmark of Nanomotif and Modkit motif identification performance. Running modkit was infeasible for the soil sample and is hence not included. Total time benchmark was performed using 144 AMD EPYC 7H12 CPUs.

| *Dataset* | *Tool* | *CPU Time [h]* | *Total Time [h:m:s]* | *Time difference [fold]* |
| --- | --- | --- | --- | --- |
| *Fecal, simple* | modkit | 6.7 | 6:32:03 | 120x |
|  | nanomotif | 0.8 | 0:03:14 | 1x |
| *Fecal, complex* | modkit | 18.0 | 1 day, 2:11:43 | 35x |
|  | nanomotif | 17.0 | 0:44:15 | 1x |
| *ZymoFecal* | modkit | 14.0 | 14:28:34 | 38x |
|  | nanomotif | 7.6 | 0:22:17 | 1x |
| *Anaerobic Digester* | modkit | 86.0 | 1 day, 23:51:30 | 23x |
|  | nanomotif | 65.0 | 2:02:50 | 1x |

## Supplementary Note 1

### Direct motif identification in contigs

The assembly sequence and the methylation pileup from “modkit pileup” are used to identify methylated motifs.

Motifs are identified in each contig sequence separately from other contigs in an assembly. We use the “fraction modified” value in the modkit pileup output to determine if a position on the contig is methylated. “Fraction modified” corresponds to the number of mapped reads modified at the position divided by the number of valid bases at the position, which is the number of reads with the same canonical base as the respective modification type (A for 6mA and C for 5mC/4mC).

Each position with a valid base is classified as; NA, if the coverage below the threshold (default 5), not methylated, if the fraction of read methylation is below threshold (default 0.7) or methylated. We further define two ways in which a position can be methylated; generally methylated positions, where fraction of read methylation is above the threshold (default 0.7) and confidently methylated position, where fraction of read methylation is above the threshold (default 0.8).

Motif search is initiated at a seed motif (the default is the respective base to the evaluated methylation type, C for 5mC & 4mC and A for 6mA). To determine which position to expand from the motif, we extract sequences in a window around all confidently methylated positions (default window size is 41, 20 bases upstream and 20 bases downstream of the methylated position). These sequences are aligned with respect to the methylation, generating a methylated sequence set, S. A positional probability matrix (PPM) is then calculated from the set S.

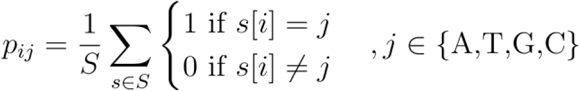

This generates a 4×41 table, where the 41 columns correspond to the relative position with respect to the methylation and the 4 rows correspond to the nucleotide. Next, 10,000 sequences of the same window size are sampled with replacement from the contig, S_sample_, and a positional nucleotide frequency table of the same dimensions is calculated. The 10,000 sampled windows serve only to generate a background positional frequency table. Resampling only reinforces the true nucleotide frequency of the background, hence resampling is non problematic.

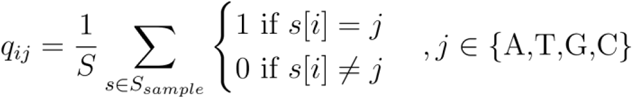

For each relative position, i, the KL-divergence is calculated from the four frequencies of the methylated sequence frequency table to the four frequencies of the sampled sequence frequency table.

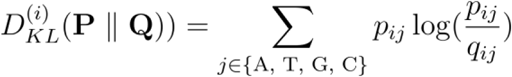

This generates a vector of size 41, where each entry corresponds to a KL-divergence value. Positions are, per default, only considered for expansion if the KL-divergence is greater than 0.05. After selecting which position to expand, we select which bases to incorporate at each of these positions by two criteria; 1. the frequency of a base in the methylation sequence frequency table must be above 25% and the frequency of a base must be above the frequency in the sampled sequence frequency table. If more than one base at a position meets this criteria, we keep both of them and combinations of them a, e.g. accepting A and G at relative position 2 with seed NNANN would give rise to NNANA, NNANG and NNANR.

Each new motif candidate after the expansion is evaluated using a beta-Bernoulli model, treating each motif occurrence as a Bernoulli trial, being a success if it is a generally methylated position and a failure if not a methylated position. We use a Beta(𝛼=0, β=1) as a prior, which means the posterior is also a Beta distribution with the parameters:

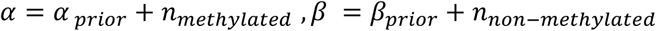

The posterior distribution is used to score each motif using the mean, standard deviation, and mean difference from the preceding motif. The mean represents the degree of motif methylation, a value expected to increase as the motif is refined. The standard deviation is used to penalize when few observations are present. Mean difference is expected to be high, when a desirable nucleotide addition is made, as it keeps the N highly methylated motif variants and disregards 4-N non-methylated motif variants, and is approximately zero for nucleotide insertion which contributes nothing to the recognition sequence.

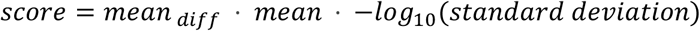

After scoring each of the new motifs, the highest scoring motif is stored. Next, one of the motifs is selected for propagation to the new set of motifs. The objective of the search is to converge on the motif candidate contributing the most positive methylation sites. The search heuristic is therefore formulated to minimize the proportion of generally methylated positions removed and maximize the proportion of non-methylated positions removed with respect to the seed motif. Concretely, the heuristic is calculated using the 𝛼 and β parameters of the beta-Bernoulli posterior of the current motif and the seed motif, as they represent the number of methylated and non-methylated motif sites.

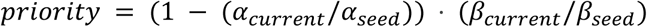

The formulation of the priority results in the motif with the lowest priority score to be chosen for the next iteration, as this decreases non-methylated sites while retaining maximum methylated sites. For the next iteration, the methylation sequences extracted initially are subsetted to those only containing the motif picked for expansion. After this the positional frequency table and KL-divergence is recalculated and the same procedure as before follows. The algorithm expands and scores following the steps described above, until the maximum score of a motif has not increased for 10 rounds or no more motif candidates are left to explore. The best scoring motif is then kept and saved to candidate motifs if its score is >0.2, otherwise dropped. The whole procedure is then repeated from the same seed, but removing sequences containing previously identified candidate motifs from methylated sequences. This is continued until 25 candidate motifs with insufficient score have been dropped or only 1% of methylation sequences remain.

After all candidate motifs have been identified in a contig, they are subjected to a series of post-processing steps to improve final motifs. First, motifs which are a sub motif of other motifs are removed, which is the case if the sequence of any other motif is contained within the sequence of the current motif, e.g. C**5mC**WGG would give rise to removal of **6mA**CCWGG, as CCWGG is contained within ACCWGG. This step was added to mitigate false positive motifs resulting from 5mC methylations in close proximity to adenine can result in 6mA methylation calls, which subsequently produce a sufficiently strong signal to “detect” 6mA motifs. In this case we accept the possibility of removing similar motifs with different methylation types. Next, we remove motifs which have isolated bases, defined as a non-N position with at least 2 Ns on both sides. Next, we merge motifs whose sequences are similar, which can be the case for more generic motifs such as CCWGG, where CCAGG and CCTGG were found as separate motifs, but should constitute one motif. Motif merging is done by constructing a distance graph between all motifs, where motifs are only connected if the hamming distance is 2 or less. Motifs are then defined to be part of the same cluster in the graph if they are mutually reachable. All motifs within the same cluster are merged into a single motif, representing all motifs contained within the cluster. The merged motif is only accepted if the mean degree of methylation is not less than 0.2 of the mean methylation of the pre merge motifs, otherwise the pre-merge motifs are kept as is. Finally, motifs are queried for motif complements. If another motif is the complementary sequence of the motif, it gets removed and added as a complementary motif instead. Palindromic motifs are always considered as the complementary of itself.

### Indirect motif detection

Direct motif identification is performed on one contig without any information from other contigs in an assembly. To detect potentially missed motifs in contigs, we perform what we term indirect detection of motifs in contigs, so called as they are only detected because the motif was directly detected with high confidence in another contig. To get indirectly identified motifs, we take the complete set of all motifs identified in all contigs and calculate 𝛼 and β of the Beta posterior of the beta-Bernoulli model for all contigs. We report the 𝛼 and beta parameters as the number of motif methylations and non-methylations, respectively.

### Bin consensus

Bin consensus is evaluated by taking the complete set of motifs for a bin and checking if a motif meets a set of criteria. Firstly, a motif has to have been directly detected in at least one of the contigs in the bin. Next, we remove motifs that are not methylated in at least 75% of the contigs in the bin. We estimate this by counting the number of motif occurrences in contigs with a mean methylation of a motif above 25% and dividing by the total number of motif occurrences in the bin; if the fraction of motif occurrences present in methylated contigs is above 0.75, they are kept. Lastly, of the kept motifs, sub-motifs are removed as described in the post-processing step in the direct motif identification section. The remaining motifs are considered bin consensus motifs.

## Supplementary Note 2

### Annotation of gold standard proteins and their methylation motifs

We initially analyzed 11 prokaryotic strains known to encode gold-standard (GS) RM system proteins using Nanomotif (Fig. S9a, supplementary data 1 & 2). GS proteins are those whose biochemical functions have been experimentally validated with their exact coding DNA sequences identified^40^. The MTase-linker module of Nanomotif successfully annotated 43 out of 44 gold-standard MTase enzymes and linked the associated motifs, if active, across the 11 monocultural genomes. The single missing annotation is likely a false negative, as DefenseFinder had multiple HMM-profile hits for this gene, but ultimately assigned the corresponding gene as part of a non-methylation mediated defense system. Among the 44 GS MTases, 40 were found to be active. Notably, the motifs identified for all active GS MTases precisely match the specificities reported for the GS MTases in REBASE. For the non-GS epigenetic systems in these 11 prokaryotic strains, our gene annotations and motif assignments are generally aligned with existing REBASE entries. However, in *S. oneidensis*, *K.aquimarina,* and *D. acetoxidans*, the MTase-linker module annotated additional type II MTases beyond those previously reported in REBASE. While some may represent false positives, for example, *contig_2_3607* in *S. oneidensis*, others are supported by active motifs without any alternative gene assignments, for example *contig_1_1589* in *K. aquimarina* (supplementary data 2).

In total, 42 out of 71 detected motifs were confidently linked to annotated RM systems or orphan MTases. For 18 additional motifs, candidate genes with matching methylation features were identified. Notably, the remaining motifs include several from *Anabaena variabilis,* that may represent variants of fewer distinct motifs, and *M.ruber,* representing noise motifs attributed to the increased false positive rate of ONT’s methylation models in high GC contexts (Fig. S3). Apart from these unassigned motifs and two motifs in *D. acetoxidans*, all detected motifs were either supported by REBASE entries or corroborated by previous PacBio sequencing data^41^, further validating the accuracy and reliability of the detected motifs.

## Supplementary Note 3

### Discovery of epigenetics systems in diverse metagenomes

We next analyzed a diverse set of prokaryotic communities using nanomotif (Fig. S10). Using the MTase-linker, 3123 MTase genes were detected across 549 HQ MAGs, resulting in a median of 6 MTases genes per genome, and 3 RM systems encompassing an MTase per genome. Type II MTases were the most abundant (67%). This was followed by type I (16%), IIG (9%), and III (7%). Type II MTases were also the most prevalent of all types not associated with an RM-system. Only 24% of type II MTases were co-located with a cognate restriction enzyme. Previous studies have also reported the frequent presence of orphan MTases in a wide range of bacterial genomes^41^. However, it is important to acknowledge that some associated REase genes may have gone undetected due to sequence divergence, especially in the complex samples with high novelty. In such cases, there may be an overestimation of orphan type II MTase genes. Despite this, many examples of orphan MTases are indeed genuine, and they represent a large group of MTases with non-RM functions. Similarly, 24% and 39% of Type I and Type III MTases, respectively, were unexpectedly identified without corresponding restriction enzymes or Type IS subunits. In the soil sample, a significantly lower number of MTase genes per genome compared to the other metagenomes was observed as well. This discrepancy is likely due to the limited sensitivity of the HMM models in complex samples.

For 76% of detected motifs, at least one candidate MTase with similar methylation characteristics was found within the same genome. In 232 cases, a single motif could be confidently linked to a specific MTase gene or RM system, resulting in a high-confidence set of MTase target motif annotations. Notably, 65 of these motifs involved 5mC modifications, which are notoriously difficult to detect with SMRT sequencing. This highlights the potential of Nanomotif to accurately annotate MTase target recognition motifs in metagenomes, including those with 5mC modifications.

## Supplementary Data

**Supplementary data 1:** a) expected motifs in REBASE gold standard monocultures. b) summary stats for recovered MAGs in metagenomes. c) contigs annotated as mobile genetic elements. d) Contamination and completeness before and after Nanomotif contamination removal. e) Motifs identified with Nanomotif in monocultures. f) Motifs identified with Nanomotif in Zymo reference samples. g) Motifs identified with Nanomotif in metagenomes.

**Supplementary data 2:** MTase-linker results from monocultures and metagenomic samples

